# Transcriptomic and functional profiling reveal autophagy inhibition and persistent bioenergetic collapse following parallel photodamage to lysosomes and mitochondria

**DOI:** 10.1101/2025.08.21.671577

**Authors:** Márcia Silvana Freire Franco, Felipe Gustavo Ravagnani, Suely Kazue Nagahashi Marie, Sueli Mieko Oba-Shinjo, Leonardo Vinicius Monteiro de Assis, Maurício S. Baptista

## Abstract

Photodynamic therapy (PDT) using 1,9-dimethyl methylene blue (DMMB) induces coordinated mitochondrial and lysosomal damage and results in strong cellular death induction. However, the underlying transcriptional regulation in response to DMMB remains elusive. We compared the transcriptome response of photoactivated DMMB (paDMMB) to the gene signature triggered by autophagy-modulating agents: rapamycin (an autophagy activator) and bafilomycin A1 (a lysosomal acidification inhibitor).

Transcriptome analysis revealed a pronounced transcriptomic response to paDMMB, with 884 differentially expressed genes (DEGs), compared to 291 for bafilomycin and 154 for rapamycin. paDMMB treatment upregulated genes associated with autophagy, mitochondrial stress responses, and proteostasis, while downregulating genes involved in miRNA processing and lipid catabolism. Rapamycin treatment downregulated amino acid biosynthesis pathways, while upregulating processes associated with nutrient starvation. Conversely, bafilomycin treatment upregulated genes related to lipid metabolism, while suppressing cytoskeletal programs. Transcriptomic comparisons revealed a striking overlap (95%) between paDMMB and bafilomycin signatures.

Among the several biological processes affected by paDMMB, mitochondrial-related processes were strongly enriched. To determine whether the acute transcriptome changes caused by paDMMB led to persistent functional effects, we stimulated cells with DMMB and assessed mitochondrial respiration after a recovery period. paDMMB reduced basal respiration, ATP production, proton leak, and maximal respiration. These effects were not further altered by bafilomycin co-treatment but were markedly exacerbated by rapamycin.

Collectively, we show that paDMMB leads to a transcriptome rewiring, closely resembling autophagy inhibition with a sustained mitochondrial dysfunction. These findings provide a valuable resource to understand the interplay between DMMB-induced lysosomal stress, transcriptional regulation, and PDT.

## INTRODUCTION

Macroautophagy is a lysosome-dependent degradation process that starts with the generation of phagophores and then evolves into autophagosomes. The fusion of autophagosomes with lysosomes produces autolysosomes, where cellular components are degraded and recycled to support energy and nutrient balance^1^. While autophagy typically serves a protective role, it can also contribute to cell death^2^. A selective form of autophagy, mitophagy, eliminates damaged mitochondria and can prevent apoptosis by preserving mitochondrial integrity ^3^.

Photodynamic therapy (PDT) employs light-sensitive photosensitizers to produce reactive oxygen species (ROS) that lead to oxidative stress across various organelles. Such responses result in the upregulation of antioxidant defenses like glutathione and glutathione peroxidase 4 (GPX4) ^4^, as well as heat shock proteins (HSPs) that stabilize misfolded proteins ^5^. PDT has been demonstrated to initiate autophagy by activating transcription factors like HIF-1α, NRF2, p53, and FoxO3 ^6,7^. Clearance of damaged peroxisomes and mitochondria, via pexophagy and mitophagy, respectively, plays a critical role in maintaining cellular health during oxidative stress ^8^.

We recently demonstrated that 1,9-dimethyl methylene blue (DMMB), a phenothiazinium-based PDT photosensitizer, achieves exceptional cytotoxicity under remarkably mild conditions (IC50 ≈ 10 nM in 10^5^ cells with only 5 min irradiation). This potency exceeds conventional PDT photosensitizers by two orders of magnitude, as typical agents require micromolar concentrations to induce comparable damage in vitro. We attribute this dramatic enhancement in cell death efficiency to DMMB’s unique capacity to simultaneously target mitochondria and lysosomes, activating regulated cell death pathways through selective organelle photodamage ^9^. Towards a better understanding of the role of autophagy in DMMB-mediated phototoxicity, we analyzed the transcriptome signature of photoactivated DMMB and compared it with the signatures of cells undergoing autophagy or autophagy inhibition. Specifically, we employed bafilomycin A1 (a V-ATPase inhibitor that blocks autophagic flux by impairing lysosomal acidification, causing lysosomal swelling, ER stress, and mitochondrial dysfunction; ^10,11,11^ and rapamycin, which induces autophagy through mTORC1 inhibition, promoting metabolic adaptation and stress resilience ^12^.

In this study, comparative transcriptomic profiling revealed distinct molecular programs and pathways linked to autophagy regulation, lysosomal stress, and mitochondrial dysfunction. Functional analyses focusing on mitochondrial respiration revealed that DMMB effects lead to sustained mitochondrial respiration impairment, following DMMB exposure. Our findings integrate molecular and functional assessment to uncover the molecular mechanisms driving DMMB-induced phototoxicity.

### Overview of the transcriptomic response to photoactivated DMMB compared to autophagy modulators

To gain a thorough understanding of the molecular mechanisms involved in DMMB treatment, we loaded human nonmalignant immortalized keratinocytes (HaCaT) cells with DMMB (10 nM) and subsequently stimulated them with red light to photoactivate DMMB (paDMMB). We also exposed HaCaT cells to Rapamycin (100 nM) and Bafilomycin (5 nM), which are known to activate and inhibit autophagy, respectively ^13,14^ to gain molecular insights into paDMMB-induced autophagy modulation. This framework served as a basis to investigate the possible effects of paDMMB in autophagy modulation (Figure 1A).

**Figure 1.**
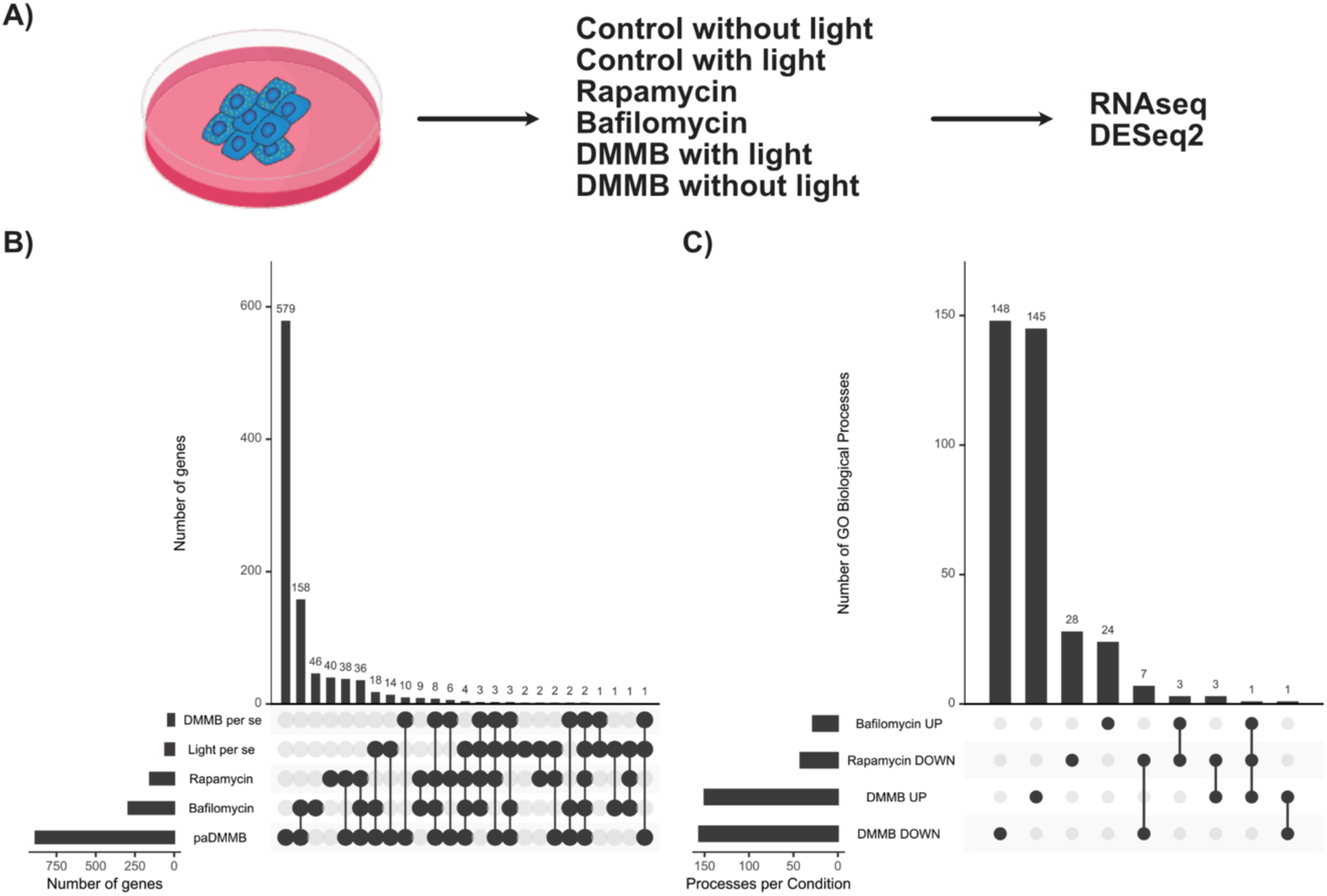
Overview of the transcriptome alteration in HaCat cells in response to different treatments. A) The experimental protocol is illustrated. B) The UpSet plot shows the count of differentially expressed genes (DEGs) detected under each condition using the Likelihood Method (LRT) in DESeq2 (padj < 0.05) compared to the control group. C) The UpSet plot displays the number of enriched biological processes associated with the DEGs identified in panel B. The image depicted in A was obtained from Bioicons.

In our analysis, we accounted for the individual effects of each factor (e.g., photoactivation, DMMB, rapamycin and bafilomycin) using the Likelihood Ratio Test (LRT) method in DESeq2 ^15^. The LRT method evaluates a complete model encompassing all experimental factors against a simplified model that includes only the intercept, which assumes no impact from any factors. Compared to the unexposed control group (padj < 0.05), the treatments resulted in distinct sets of differentially expressed genes (DEGs): 99 for rapamycin, 291 for bafilomycin, and 884 for paDMMB. The effect of DMMB in the absence or in the presence of photoactivation protocols led to a marginal response of 39 and 57 DEGs, respectively. However, only two genes (e.g., TBL1X and S100A8) were exclusively identified as being influenced by the photoactivation protocol (Table S1).

Given the minimal transcriptional effect of DMMB alone and light exposure per se, we focused subsequent analyses on the main treatment groups: rapamycin, bafilomycin, and paDMMB. Most DEGs were unique to each condition, with limited overlap across treatments (Figure 1B). Gene ontology enrichment analysis of these DEGs revealed distinct biological processes associated with each treatment, showing the condition-specific nature of the transcriptional responses (Figure 1C).

### Overview of the transcriptome signature evoked by rapamycin

In order to investigate the transcriptomic changes induced by rapamycin, we focused on genes associated with rapamycin treatment (Figure 2A), a well-known autophagy inducer ^13,16–18^. A total of 154 DEGs were identified in the rapamycin-treated group (82 downregulated and 72 upregulated genes, padj < 0.05 and log2FC ± 0.58). Enrichment analysis of these DEGs confirmed the metabolic effects caused by rapamycin. Downregulated genes were enriched for several metabolic pathways, including amino acid metabolism, monocarboxylic acid and fatty acid metabolism, carbohydrate metabolism, and nutrient sensing. Conversely, enrichment analysis of the upregulated genes identified processes related to lipid and triglyceride homeostasis, response to starvation, and inflammatory signaling. Additionally, pathways related to angiogenesis, endothelial cell migration, NF-κB signaling, and interleukin-6 production were overrepresented (Figure S1; Table 1).

**Figure 2.**
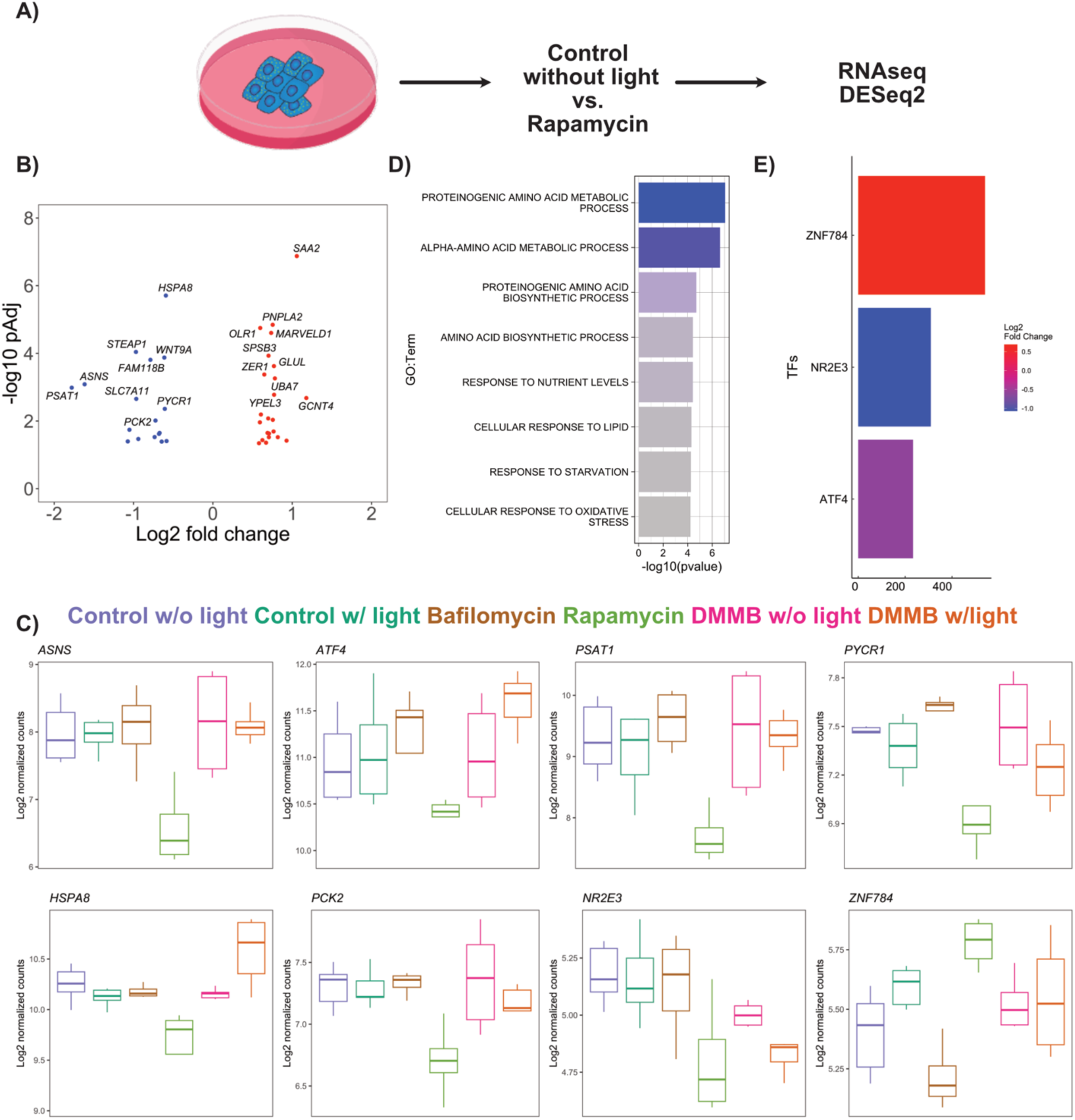
Transcriptome signatures in response to rapamycin treatment. A) The experimental setup is illustrated. B) The volcano plot displays the unique DEGs (identified solely in the Rapamycin-treated group). C) Representative unique Rapamycin DEGs are displayed. D) Enrichment analyses conducted for downregulated genes (in blue). E) Predictive transcription factors based on the unique DEGs (shown in B) are presented. The mean score is represented on the x-axis and the log2 fold change on the y-axis. The Image described in A was obtained from Bioicons.

We focused on identifying specific genes associated with Rapamycin by filtering our dataset to find exclusive DEGs. A total of 40 DEGs (17 downregulated and 23 upregulated genes, padj < 0.05 and log2FC ± 0.58) were exclusively identified in response to rapamycin treatment (Figure 2B; Table S1). Representative DEGs are shown in Figure 2C. No enriched pathway was identified for the upregulated DEGs. In contrast, the downregulated genes were significantly enriched in multiple amino acid metabolic and biosynthetic processes, reflecting the metabolic impact of the treatment ^17^ (Figure 2D; Table S1). Specifically, processes such as proteinogenic amino acid metabolism, L-amino acid metabolism, and alpha-amino acid metabolism were highly represented (e.g., *ASNS*, *ATF4*, *PSAT1*, *SLC7A11*, and *PYCR1*). Additionally, genes involved in the glutamine family amino acid metabolic process and amino acid biosynthetic process also showed a downregulation (*ASNS*, *SLC7A11*, and *PYCR1*), further suggesting suppression of essential components of protein and nitrogen metabolism. A total of three predicted transcriptional factors, such as *ATF4*, *NR2E3*, and *ZNF784*, were identified (Figure 2C and E).

Overall, our annotated metabolic processes showed a broad suppression of amino acid turnover and biosynthesis, reflecting reduced anabolic activity, in accordance with well-known effect of rapamycin in the inhibition of the mTORC1 pathway, a key nutrient sensor regulating protein synthesis and metabolism^17^. Furthermore, these processes highlight rapamycin’s role in mimicking nutrient deprivation by inhibiting mTORC1, effectively triggering adaptive cellular responses that promote autophagy ^16^.

### Overview of the transcriptome signature evoked by bafilomycin

In order to investigate the transcriptomic changes induced by bafilomycin, we focused on genes associated with bafilomycin treatment (Figure 3A). Bafilomycin A1 inhibits autophagic flux in vitro by preventing lysosomal acidification. This macrolide targets the V-ATPase ATP6V0C, inhibiting protons from entering the lysosomal lumen. This disruption alters the acidic environment needed for effective lysosomal function and autophagic degradation ^14,19^. A total of 291 DEGs were identified in the bafilomycin-treated group (113 downregulated and 178 upregulated genes, padj < 0.05 and log2FC ± 0.58, Table S1). Enrichment analysis of these DEGs confirmed the metabolic effects caused by bafilomycin. Downregulated genes enriched for development, cytoskeleton, migration as well as metabolic processes (e.g., lipid and steroid metabolism). Whereas upregulated genes were enriched for processes associated with lipids, cholesterol, and carbohydrate metabolism, inflammatory and wound healing response, hormone secretion, apoptosis and extracellular matrix organization (Figure S2; Table S1).

**Figure 3.**
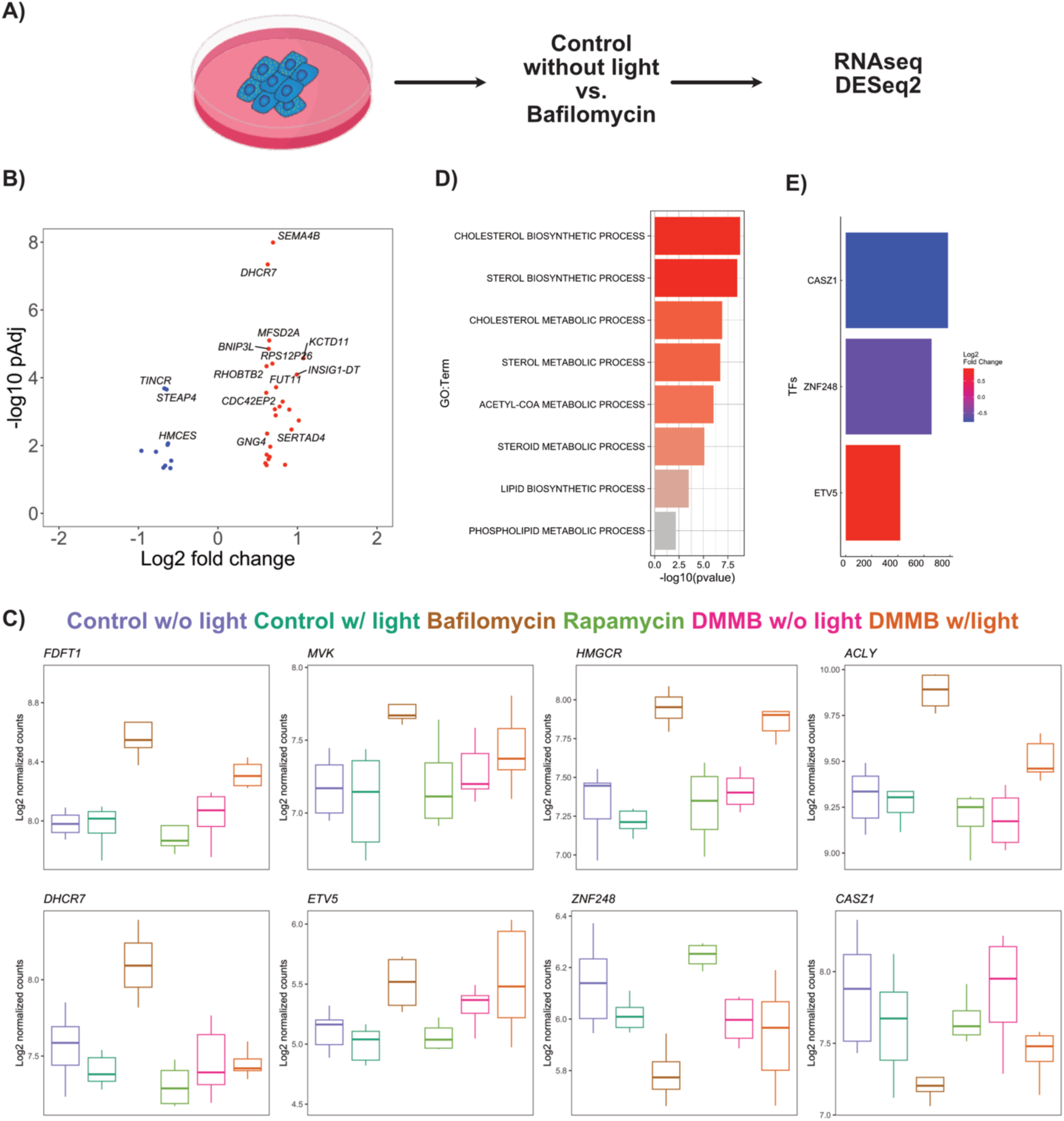
Transcriptome signatures in response to bafilomycin treatment. A) The experimental setup is illustrated. B) The volcano plot displays the unique DEGs (identified solely in the bafilomycin-treated group). C) Representative unique bafilomycin DEGs are displayed. D) Enrichment analyses conducted for upregulated genes (in red). E) Predictive transcription factors based on the unique DEGs (shown in B) are presented. The mean score is represented on the x-axis and the log2 fold change on the y-axis. The Image described in A was obtained from Bioicons.

Focusing on identifying specific genes associated with bafilomycin, we filtered our dataset to find exclusive DEGs. A total of 46 DEGs (10 downregulated and 36 upregulated genes, padj < 0.05 and log2FC ± 0.58) were exclusively identified in response to bafilomycin treatment (Figure 3B; Table S1). Representative DEGs are shown in Figure 3C. While no pathways were enriched among downregulated DEGs, upregulated genes were significantly associated with lipid and cholesterol metabolism, alcohol and isoprenoid pathways, acyl-CoA and pyruvate metabolism, and small molecule biosynthetic processes (Figure 3D; Table S1). A total of three predicted transcriptional factors (TFs), such as *ETV5*, *ZNF248*, and *CASZ1*, were identified (Figure 3C and 3E).

Overall, the pathways triggered by bafilomycin treatment indicate a metabolic shift affecting both carbohydrate and lipid metabolism because of bafilomycin’s role as a lysosomal acidification inhibitor and autophagic influx. These effects on autophagic influx led to a compensatory, transcriptional response in several energy-related pathways.

### Overview of the transcriptome signature evoked by paDMMB

Previously, we identified that at low concentrations (10 nM), paDMMB induced mitochondrial damage and mitophagy, but lysosomal damage prevented its completion, leading to enhanced cell death. This parallel damage strategy led to a significant improvement in photoinduced cell death ^9^.

To gain a more comprehensive understanding of the impacted transcriptional programs, we examined the transcriptome of HaCaT cells treated with paDMMB alone (Figure 4A). A total of 884 DEGs were identified in the paDMMB-treated group (343 downregulated and 541 upregulated genes, padj < 0.05 and log2FC ± 0.58, Table S1). Enrichment analysis of these DEGs showed a strong impact on the transcriptome. For example, downregulated DEGs were linked to processes involving development and differentiation, miRNA metabolism, lipid breakdown, and signal transduction (Notch, SMAD, and MAPK signaling). Conversely, upregulated DEGs were associated with inflammation and immune response, cell migration, apoptosis, protein folding, lipid/cholesterol metabolism, autophagy, and proteostasis (Figure S3; Table S1).

**Figure 4.**
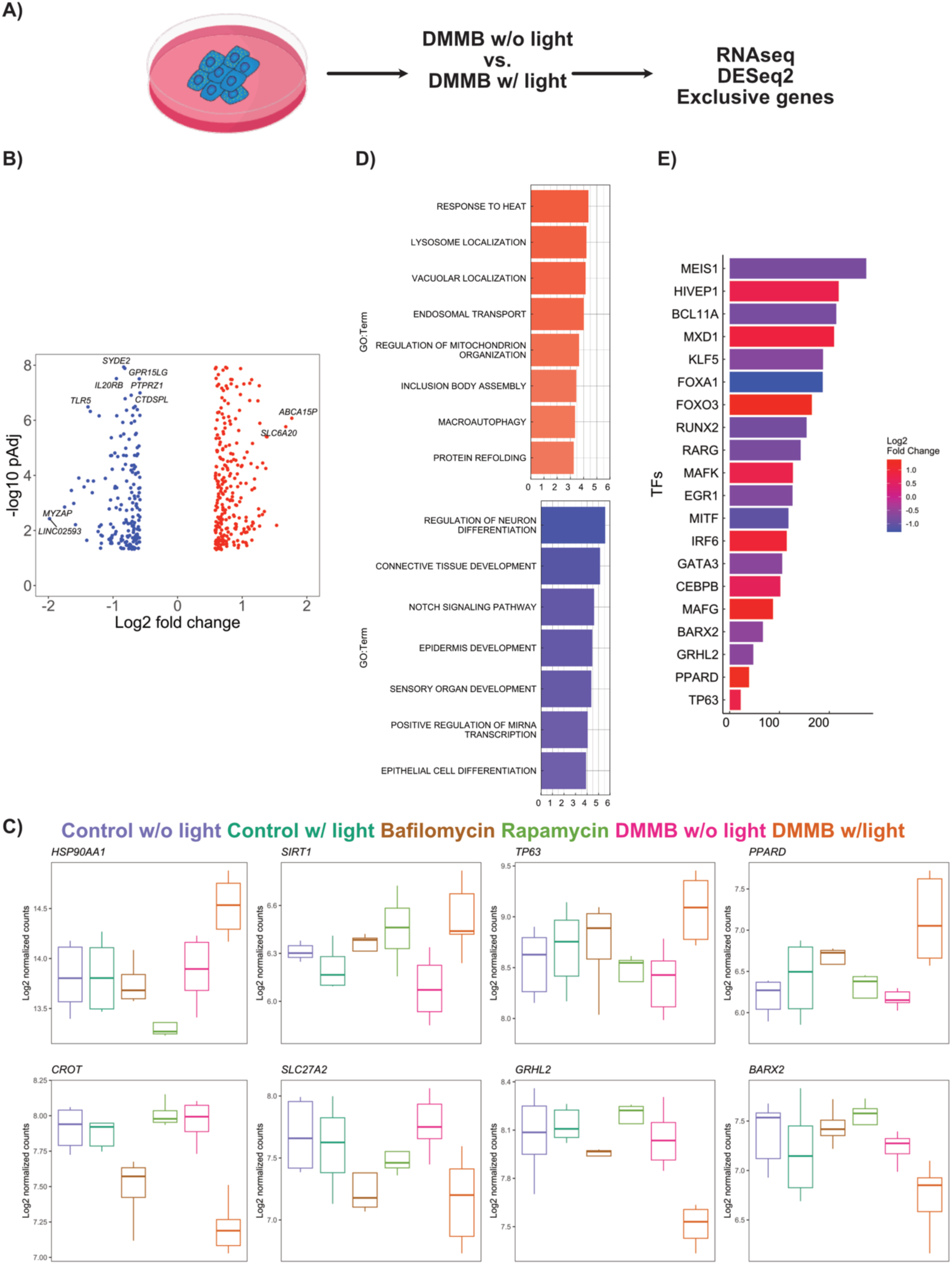
Transcriptome signatures of photoactivated DMMB. The experimental setup is illustrated. B) The volcano plot displays the unique DEGs (identified solely in the DMMB). C) Representative unique DMMB DEGs are displayed, with the first and second rows showing upregulated and downregulated DEGs, respectively. D) Enrichment analyses conducted for upregulated genes (in red) and downregulated genes (in blue). E) Predictive transcription factors based on the unique DEGs (shown in B) are presented. The mean score is represented on the x-axis and the log2 fold change on the y-axis. The Image described in A was obtained from Bioicons.

Focusing on identifying specific genes associated with paDMMB, we filtered our dataset to find exclusive DEGs. A total of 579 DEGs (215 downregulated and 364 upregulated genes, padj < 0.05 and log2FC ± 0.58) were exclusively identified in response to paDMMB treatment (Figure 3B; Table S1). Representative DEGs are shown in Figure 4C. Exclusively downregulated genes in response to paDMMB were significantly enriched for processes related to development, differentiation, and stem cell fate determination, together with pathways involved in post-transcriptional and post-translational regulation (e.g., miRNA transcription). In contrast, exclusively upregulated genes were strongly associated with cellular stress response and homeostasis, including apoptosis regulation, autophagy, and proteostasis. Key pathways included the regulation of mitochondrial dynamics (e.g., fission, membrane potential, and organelle organization) and immune-related signaling (e.g., cytokine production and inflammation). Additional enrichment was observed in genes associated with proteolysis and ubiquitin-mediated protein degradation, as well as responses to unfolded or misfolded proteins (Figure 4D; Table S1). TF prediction analyses identified a total of 49 genes, such as *TP63*, *PPARD*, *GRHL2*, *BARX2*, *MAFG*, *CEBPB*, and *GATA3* (Figure 4C and E).

Overall, the transcriptional response to paDMMB is characterized by a strong upregulation of stress-related programs, including autophagy, mitochondrial regulation, proteolysis, and apoptosis, accompanied by a marked downregulation of developmental pathways linked to cell differentiation, stem cell fate, and post-transcriptional regulation.

### Transcriptome signature comparisons reveal a strong link between the paDMMB-induced and the bafilomycin-induced signatures

Given the pronounced transcriptional changes induced by paDMMB treatment, we analyzed the paDMMB-induced transcriptional signature and compared it to the modifications elicited by autophagy modulation, specifically through either the induction with rapamycin or the inhibition with bafilomycin. We hypothesized that if the transcriptional profiles of paDMMB-treated cells are further shaped in a direction similar to known drug-specific signatures, the resulting overlap in gene expression could suggest how autophagy contributes to the effects of paDMMB.

We found that only a limited number of genes (46 out of 99 DEGs) showed consistent direction of change (e.g., UP in paDMMB and UP in the rapamycin group) between the transcriptomes of paDMMB and rapamycin-treated cells (Figure 5A – B; Table S1). In contrast, we observed a significant overlap of 95% (221 out of 231 DEGs) with the same direction of change between the paDMMB and bafilomycin groups, reflecting comparable enriched processes identified between these conditions (Figure 5 C – D; Table S1).

**Figure 5.**
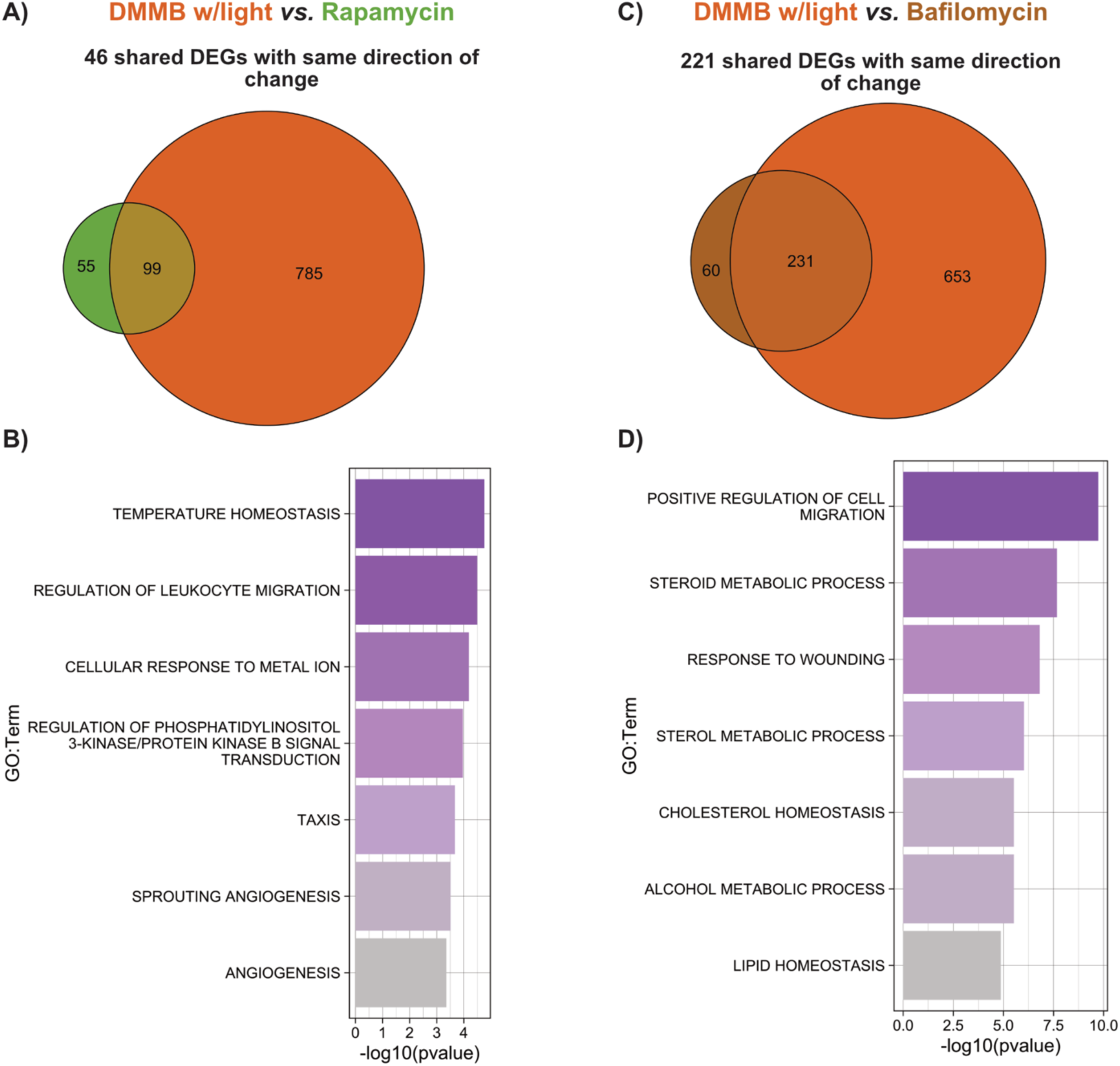
Comparison of transcriptome signatures between DMMB and Rapamycin or Bafilomycin treated groups. A – C) Venn diagram depicts the shared differentially expressed genes (DEGs) between DMMB and Rapamycin (A) or Bafilomycin (C). B – D) Enrichment analyses of the shared DEGs with the same regulation identified for each comparison.

Overall, our analysis suggests that the transcriptional effects of paDMMB resemble an autophagy inhibition profile, as evidenced by the high DEGs overlap and similar enriched biological processes observed with bafilomycin treatment. Conversely, the activation of autophagy with rapamycin via rapamycin does not yield a similar impact, indicating that the molecular outcomes of paDMMB are more closely associated with pathways disrupted by lysosomal inhibition than by mTOR suppression.

### Mitochondrial Respiration in response to DMMB treatment

One of the most impacted processes in paDMMB was those related to mitochondria (Figure 4D). While transcriptome profiling offered a detailed view of the immediate effects following paDMMB within 6 hours, it only captured short-term changes. To assess whether these alterations led to lasting mitochondrial dysfunction, we conducted a Seahorse assay to evaluate mitochondrial function. Right after irradiation, cells were cultured in a mild nutrient– and growth-factor–restricted medium (1% FBS) for 12 hours, to enhance the cellular response during early recovery. Then, they were switched back to a nutrient-rich medium (10% FBS) for 36 hours before Seahorse analysis. This longer period helped us determine if mitochondrial impairments persisted beyond the initial injury, allowing us to distinguish between transient phototoxic effects and sustained deficits. Additionally, we tested the impact of paDMMB in the presence or absence of autophagy modulators like rapamycin and Bafilomycin.

Mitochondrial function was assessed by real-time monitoring of the oxygen consumption rate (OCR; primarily consumed by glucose metabolism through the citric acid cycle). Figure 6A shows a typical experimental scheme in which specific inhibitors were added to respiratory complexes, allowing for the evaluation of various respiratory chain components, including basal respiration, ATP-linked respiration, proton leak, and maximal respiration.

**Figure 6.**
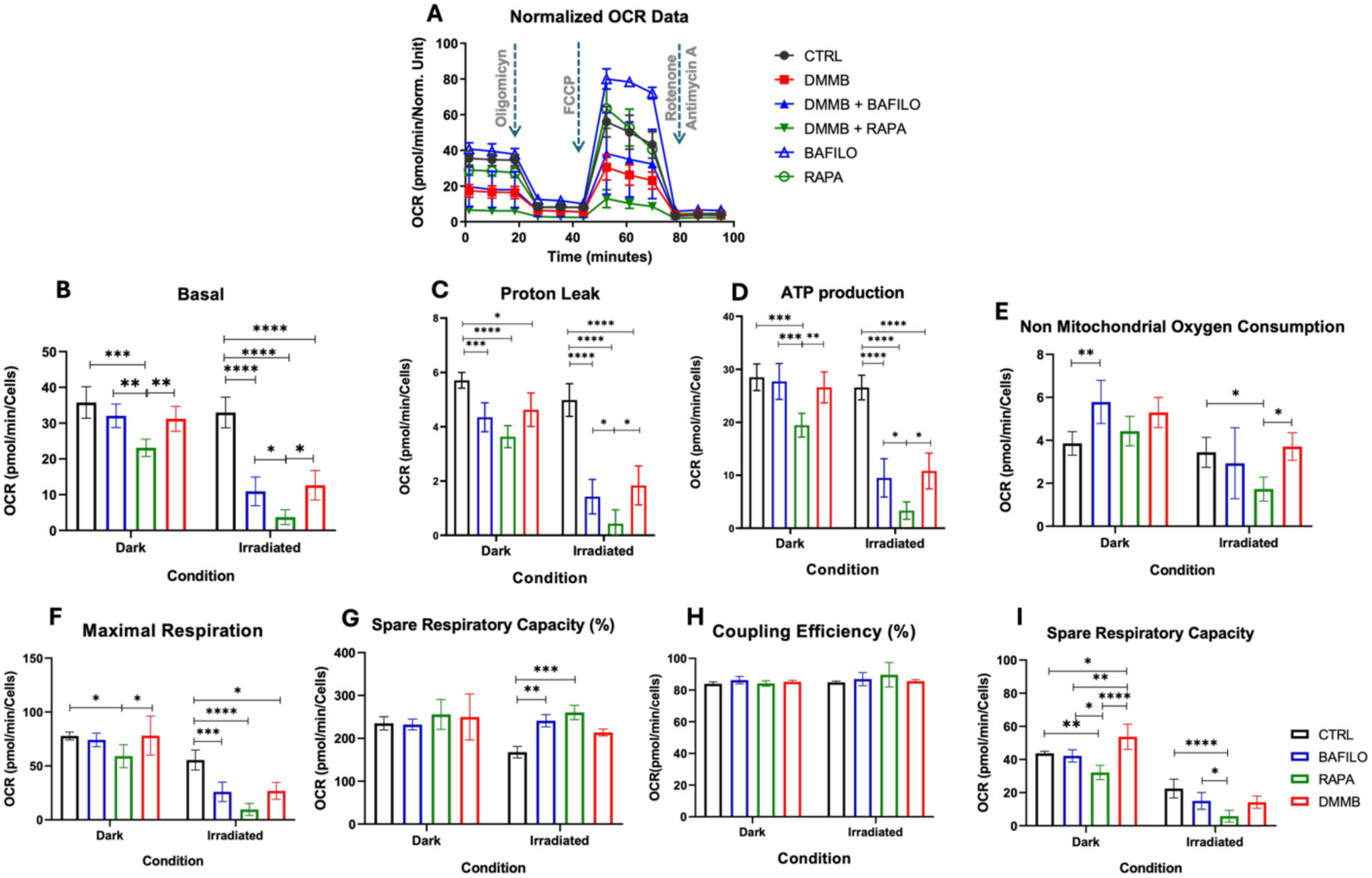
Mitochondrial response in response to DMMB in the presence of rapamycin or bafilomycin. Oxygen consumption rate (OCR) from HaCat cells exposed to DMMB (10 nM), Bafilomycin (BAFILO), Rapamycin (RAPA), DMMB+BAFILO, and DMMB+RAPA for 24 h was determined by the mitochondrial stress test. OCR was measured before and after sequential injection of Oligomycin (0.1 µg/mL), FCCP (1 µM), and Antimycin A (10 µM). A) mitochondrial respiration by Seahorse, following the mitochondrial stress analysis. The oxygen consumption rate (OCR) curves along the time interval up to 60 min are presented according to applied drugs. **B**) Bar chart of basal respiration; C) Proton Leak; D) ATP production; E) Non-Mitochondrial Oxygen Consumption; F) Maximal respiration; G) Spare Respiration Capacity; H) Coupling Efficiency; I) Spare Respiratory Capacity. * p< 0.05, ** p< 0.01, *** p< 0.001, **** p< 0.0001.

Cells treated with Bafilomycin or DMMB in the dark group (not exposed to red light) had no significant differences in basal respiration (Figure 6B), ATP production (Figure 6D), and maximal respiration (Figure 6F) when compared to the control. The exception was a slight increase in spare respiratory capacity in the group that received DMMB without photoactivation (Figure 6I). The effects of bafilomycin in the absence of paDMMB resulted in higher non-mitochondrial oxygen consumption (Figure 6E). However, rapamycin-treated cells in the absence of paDMMB showed a decrease in basal respiration (Figure 6B), proton leak (Figure 6C), ATP production (Figure 6D), maximal respiration (Figure 6F), and spare capacity (Figure 6I). These results clearly show a strong effect of rapamycin on mitochondrial respiration.

In the presence of paDMMB, the cells showed reduced basal respiratory (Figure 6B), proton leak (Figure 6C), ATP production (Figure 6D), and maximal respiration (Figure 6F) when compared to the control group. Importantly, the effects of DMMB were similar to those evoked in the BAFILO-DMMB group, demonstrating no additional effects. However, the effects of DMMB were maximized in the presence of rapamycin, leading to further decreases in basal respiration (Figure 6B), proton leak (Figure 6C), ATP production (Figure 6D), and non-mitochondrial oxygen consumption (Figure 6E).

Overall, paDMMB caused widespread and lasting impairments in mitochondrial respiration, an effect that was significantly worsened by rapamycin but not further changed by bafilomycin.

## DISCUSSION

### Rapamycin effects

The downregulation of several metabolic processes associated with amino acid, lipid, and fatty acid metabolism and biosynthesis reflects a shift away from anabolic processes, aligning with autophagy activation. In contrast, the upregulation of pathways involving acylglycerol and triglyceride, starvation, and chemokine (e.g., IL-6) suggests an adaptive mechanism to cope with nutrient and energy stress, reinforcing its impact on autophagic and metabolic regulation ^20^. This shift is orchestrated by central metabolic regulators such as the mammalian target of rapamycin (mTOR) and AMP-activated protein kinase (AMPK), which integrate signals from amino acid availability, energy status, hormone and growth factor signaling, and various stress conditions (e.g., oxidative stress, hypoxia, DNA damage). AMPK and mTORC1 have an antagonistic relationship: whereas mTOR stimulates anabolic processes, its inhibition promotes autophagy and enhances metabolic flexibility, supporting cellular survival and maintaining energy balance under balanced stress ^21–23^.

We observed a significant downregulation of genes related to protein synthesis and extracellular signaling, consistent with mTORC1 inhibition, and possibly AMPK activation ^20,24^. Glucose deprivation typically reduces intracellular ATP levels, decreasing the ATP/ADP and ATP/AMP ratios, which can activate AMPK. Cells treated with rapamycin had increased expression of *ZBTB20* (Zinc finger and BTB domain-containing protein 20), a transcriptional repressor involved in cellular development, metabolism, differentiation, and innate immunity ^25^. However, this gene was not exclusively associated with rapamycin effects. The role of lipid mobilization to meet energy demands is evident through the upregulation of several genes, such as *RORA*, *FGF1*, and *PNPLA2*. For instance, PNPLA2 catalyzes triglyceride hydrolysis and releases free fatty acids for mitochondrial β-oxidation during energy depletion. In the context of autophagy, PNPLA2 activity can complement lipophagy, enhancing the supply of fatty acids from lipid droplets ^26,27^.

Oxidative stress is likely a key factor in the rapamycin-induced effects. For instance, under physiological conditions, H₂O₂ is detoxified by enzymes such as catalase, peroxiredoxins, or GSH-dependent enzymes like GPX1, which convert GSH into GSSG. However, when in excessive amounts, hydrogen peroxide (H₂O₂) can produce harmful hydroxyl radicals (·OH). The enzyme GSR regenerates GSH from GSSG using NADPH as an electron donor. This antioxidant defense system is regulated by NRF2, a redox-sensitive transcription factor that coordinates cellular responses to oxidative stress ^28^. In rapamycin-treated cells, several genes associated with oxidative stress were identified, including *HMOX1*, *SLC7A11*, *BNIP3*, and *PYCR1*.

Under amino acid deprivation, ATF4 initiates transcription of genes involved in amino acid transport and biosynthesis, including asparagine synthetase (ASNS), which increases *de novo* asparagine synthesis when asparagine levels drop ^29,30^. This upregulation promotes the expression of glutamine synthetase (*GLUL*) and lysis of glutamine, thereby facilitating the continuous use of intracellular glutamine ^31^. Through this mechanism, cells can restore global protein translation disrupted by glutamine scarcity ^31,32^. In our dataset, *ATF4* was downregulated in rapamycin-treated cells, a finding that diverges from the typical nutrient-stress signature where *ATF4* is upregulated. Previous studies identified that rapamycin’s effects are mediated by *NRF2* ^33,34^. We identified two putative novel TFs (*NR2E3* and *ZNF784*) that can mediate rapamycin effects.

Seahorse metabolic profiling showed that rapamycin alone significantly diminished basal respiration, ATP production, proton leak, and spare respiratory capacity, indicating reduced mitochondrial activity. In cells with paDMMB, mitochondrial functions were severely compromised; however, the additional treatment with rapamycin worsened this dysfunction, leading to a further decline in all essential respiratory parameters beyond what paDMMB alone caused.

These results suggest that although rapamycin triggers stress-adaptive transcriptional programs, it also restricts mitochondrial plasticity, which may make cells more susceptible to oxidative stress and energy depletion when combined with paDMMB.

### Bafilomycin effects

Bafilomycin-treated cells adjust to metabolic stress in ways that maintain self-renewal, proliferation, and survival. Under nutrient-limiting conditions, such as glucose and oxygen deprivation, cells undergo the “Warburg effect,” a shift from oxidative phosphorylation to aerobic glycolysis in the pentose phosphate pathway, resulting in lactate accumulation while maintaining ATP production ^35^. Bafilomicyn sustains this uncoupled metabolism and elevated respiration without compromising viability, a response attributed to increased cytosolic Ca^2+^ level due to V-ATPase inhibition, while the mitochondrial Ca^2+^ and mitochondrial pH gradient remain moderately affected, supporting continued ATP production via oxidative phosphorylation ^11^.

Consistent with hypoxia-like adaptation, we observed a strong upregulation of *PDK1*, whose protein inhibits the pyruvate dehydrogenase complex (PDH), limiting pyruvate entry into the TCA cycle and enhancing glycolytic flux. This is a common adaptive response to hypoxia and mitochondrial stress ^36^. These pathways are transcriptionally regulated by HIF1α, which activates *PDK1*, *FUT11*, and *BNIP3L* under low-oxygen conditions ^37,38^. In our conditions, the previous genes (*PDK1*, *FUT11*, and *BNIP3L*) were upregulated exclusively in the bafilomycin-treated group. For instance, *FUT11* has been associated with hypoxic pancreatic cancer progression through PDK1 stabilization and AKT/mTOR activation, promoting cell survival ^37^.

We also observed a coordinated upregulation of genes involved in acetyl-CoA and sterol biosynthesis, including *ACLY*, *MVD*, *MVK*, *FDFT1*, and *DHCR7*, suggesting a metabolic rerouting to support lipid synthesis. Importantly, these genes were exclusively upregulated in the bafilomycin-treated group. Acetyl-CoA serves as a crucial metabolic hub, feeding into the TCA cycle and biosynthetic pathways ^39^. In the cytosol, acetyl-CoA is produced from citrate via ATP-citrate lyase (ACLY), while pantothenate kinase (PANK) initiates Coenzyme A biosynthesis, supplying the CoA necessary for acetyl-CoA formation. In mitochondria, acetyl-CoA combines with oxaloacetate to form citrate via citrate synthase; citrate is then transported to the cytosol through the citrate transporter SLC25A1 and converted back into acetyl-CoA by ACLY, supporting lipid biosynthesis ^40^. This acetyl-CoA pool fuels the mevalonate pathway to produce isoprenoids, with squalene synthase (FDFT1) and lanosterol synthase (LSS) catalyzing key steps in cholesterol biosynthesis. Cholesterol and its intermediates contribute not only to membrane structure but also to intracellular signaling, including modulation of AKT and NF-κB pathways, thereby promoting pro-survival and stress-response gene expression ^41^. Notably, FDFT1 activity also elevates intracellular squalene, a lipid antioxidant that protects against ferroptosis ^42^. Further downstream, LSS catalyzes lanosterol formation, which supports proteostasis by promoting clearance of misfolded proteins through heat shock factor 1 (HSF1) activation ^43^. Similarly, DHCR7 expression is linked to developmental processes, cellular differentiation, and apoptosis, suggesting the physiological relevance of sterol metabolism in stress response ^44,45^.

We identified three potential TFs (*ETV5*, *ZNF248*, and *CASZ1*), which are novel to the effects of bafilomycin. Previous studies suggest that *ETV5* is involved in lipid metabolism by interacting with PPAR signaling ^46^ and as a driver for cell proliferation in different cancer types ^47^. The role of *ZNF248* is still being explored, but it has been implicated in cancer progression ^48^. Lastly, *CASZ1* was previously shown to be induced by DNA damage ^49^ and as an essential activator of epidermal differentiation ^50^.

Seahorse metabolic analysis showed that bafilomycin alone did not significantly alter mitochondrial respiration, with basal respiration, ATP production, and maximal respiration comparable to control cells. A modest increase in non-mitochondrial oxygen consumption was observed, possibly reflecting minor redox changes. In contrast, photoactivated DMMB led to a marked suppression of mitochondrial functions, and this effect was not further enhanced or mitigated by bafilomycin co-treatment. Despite transcriptomic evidence of metabolic adaptation, including upregulation of glucose metabolism, sterol biosynthesis, and hypoxia-responsive genes, these changes were insufficient to rescue mitochondrial activity.

Together, these findings suggest that bafilomycin-treated cells interact in compensatory pathways to maintain metabolic flexibility but fail to counteract the dominant mitochondrial dysfunction induced by paDMMB.

### Effects of photoactivated DMMB

The photodynamic effect in the presence of DMMB triggered major pathways related to macroautophagy, mitophagy, protein folding, and oxidative stress, which is consistent with previous observations by our group showing the formation of acidic vacuoles and damage to mitochondria and lysosomes as central events in DMMB-induced cell death ^9^. Among the autophagy-related genes upregulated, we identified *C9ORF72*, *ARL8B*, *ATP6V1G1*, *STAM,* and *CHMP1B*, supporting activation of autophagic and endolysosomal compartments. Of these genes, only *C9ORF72* and *CHMP1B* were not exclusively identified in the group that received paDMMB. *C9ORF72* positively regulates autophagosome formation by activating RAB proteins, essential for membrane trafficking and autophagosome maturation ^51^. ATP6V1G1 contributes to autophagosome acidification and degradation of damaged organelles ^52^, whereas ARL8B and STAM may recruit the autophagic machinery to damaged mitochondria. The upregulation of *CHMP1B*, a component of the ESCRT-III complex, is consistent with roles in autophagosome closure and endosomal sorting ^53,54^.

We also observed activation of SMAD-specific E3 ligase SMURF1, a regulator of TFEB-dependent lysosomal biogenesis through PPP3CB ubiquitination ^55^. The phospholipase PLAA and VCP/p97 were also induced, which may contribute to lysosomal damage resolution and autophagic clearance ^56^. This lysosomal-autophagy axis is central to stress resolution, yet its disruption likely underlies the enhanced DMMB-induced cytotoxicity that we previously described ^9^.

Moreover, the genes *HSPA1A*, *HSPA1B*, *DNAJB6*, and *BAG3*, which are involved in heat shock responses and chaperone-assisted selective autophagy, were exclusively identified in cells receiving paDMMB. Notably, *BAG3* interacts with p62/SQSTM1 to facilitate the degradation of damaged proteins and organelles. Our transcriptome data revealed overexpression of *DNAJBa* in the paDMMB group, and the upregulation of this gene has been implicated in both autophagy modulation and oxidative stress responses, promoting ferroptosis and altering mitochondrial dynamics ^57,58^. The ROS stress system activation was also enriched in cells that received paDMMB, as genes involved in oxidative stress, such as *RAC2* ^59^, *GCLC/GCLM* ^60^, and *PINK1* ^61^ were identified. This reflects the high oxidative load induced by paDMMB and the cell’s attempt to counteract it through redox balancing mechanisms.

We also noted metabolic rewiring in response to paDMMB. For example, genes related to insulin signaling (e.g., *GRB10*, *SIRT1*, *SOCS3*, *INSIG1*, *LPIN1*) were upregulated in cells treated with paDMMB. Several regulators of lipid metabolism (e.g., *SREBF1*, *EGR1*, *CROT*, *SLC27A2*, *FABP5*, *PLD1*, and *ABCG1*) were upregulated, which suggests a strong modulation of lipid metabolism in response to DMMB activation and is likely associated with autophagosome membrane dynamics ^62,63^. Interestingly, *PPARD* was upregulated, which is known to promote fatty acid oxidation and autophagy via AMPK-mTOR signaling^64^. Importantly, enrichment for insulin and lipid metabolism was not observed in genes exclusively identified in the DMMB group. Additionally, we observed the enrichment of circadian-related genes (*CRY2*, *BMAL1*, *RORA*, *BHLHE40*, *CSNK1E*) in response to paDMMB, indicating a change in circadian rhythms following DMMB activation, a common feature of stress response ^65–67^.

Finally, mitochondrial stress was evidenced by a regulation of several genes (e.g., *PINK1*, *BNIP3*, *PPIF*, GCLC, *PLD6*), which regulate mitophagy and antioxidant responses ^68^. For instance, PPIF, associated with mitochondrial permeability transition pore (mPTP), promotes membrane depolarization and interacts with PINK1 to initiate mitophagy ^69^. GCLC supports glutathione synthesis and mitochondrial redox buffering ^70^. The solute carrier family members SLC25A33 and SLC25A39, which mediate nucleotide and glutathione transport across the mitochondrial membrane, were also upregulated, suggesting sustained mitochondrial metabolism under stress ^71,72^.

These transcriptional changes were functionally supported by Seahorse metabolic flux analysis, designed to evaluate the long-term consequences of paDMMB treatment on mitochondrial function. This experimental approach allowed us to assess whether the short-term transcriptomic changes observed within 6 hours translated into sustained mitochondrial dysfunction. Specifically, DMMB treatment led to significant decreases in basal respiration, proton leak, ATP production, and maximal respiration, consistent with mitochondrial dysfunction. DMMB without photoactivation caused only minor changes, with a slight increase in spare respiratory capacity. These findings confirm that photoactivation is essential for DMMB toxicity and suggest that DMMB impairs mitochondrial function through oxidative and lysosomal stress. Importantly, blue and visible light may act as a chronic environmental stressor that gradually disrupts cellular homeostasis, triggering oxidative stress ^73^. We have recently shown that chronic exposure to blue light leads to widespread effects on nuclear morphology, chromatin organization, and transcriptional rewiring, ultimately promoting a pro-survival and potentially pre-malignant phenotype ^74^. In contrast, paDMMB induces acute, targeted cellular stress (e.g., mitochondria and lysosomes), leading to enhanced cellular death ^9^ and disrupting mitochondrial function (our current findings). While blue light induces sustained global damage over time ^74^, DMMB causes rapid, organelle-specific dysfunction upon activation. These findings highlight the distinct nature of chronic versus targeted photo-induced stress and their differential impact on cellular fate.

## LIMITATIONS

Our study has limitations. First, transcriptome profiling only captured an early response following DMMB photoactivation, whereas mitochondrial function was assessed after a recovery period, with the goal of evaluating long-term, rather than acute effects on cellular respiration. Second, a transcriptional signature comparison between the treatments suggests a strong overlap between paDMMB and bafilomycin. However, these comparisons are based on transcriptional similarities rather than direct comparative experiments. Furthermore, we can have similar gene expression profiles and biological effects while activating different mechanisms/systems for cell signaling.

## CONCLUSION

Building on our previous study showing that DMMB induces mitochondrial and lysosomal damage to impair autophagy and promote cell death ^9^, our current transcriptional and metabolic analyses reveal that this dual organelle stress triggers a coordinated stress response. Photoactivated DMMB upregulates mitophagy and lysosome-related genes while impairing mitochondrial respiration and ATP production, supporting mitochondrial failure as a key outcome. Transcriptome comparisons revealed strong pathway-level overlap with bafilomycin, but not rapamycin, indicating lysosomal dysfunction, rather than mTOR inhibition, as the dominant mechanism impairing autophagic flux. Together, these findings show that targeted disruption of autophagy via combined mitochondrial and lysosomal damage reprograms cellular metabolism and potentiates DMMB-based photodynamic therapy.

## MATERIAL AND METHODS

### Cell Culture

HaCaT cells were maintained in Dulbecco Modified Eagle Medium, DMEM (Sigma-Aldrich, D5648) supplemented with 10% (v:v) fetal bovine serum (FBS; Gibco™, 12657029), 100 units/mL of penicillin, 100 μg/mL of streptomycin and 250 ng/mL of amphotericin B in a 37°C incubator under a moist atmosphere of 5% carbon dioxide.

### Experimental protocol for RNAseq

Twenty-four hours after sending, HaCaT cells were incubated with DMMB (10 nM) for one hour, then washed with PBS twice. Then, irradiation was carried out with specialized equipment (Ethik, Biolambda, Brazil) with a maximum emission wavelength at 630 nm. To provide 12 J.cm^−2^, cells were irradiated for 9 min (23,5 mW). In parallel, a different batch of cells was treated with 5 nM Bafilomycin-A1 (Bafilomycin) or 100 nM Rapamycin. All samples were collected 6h later. Total RNA was extracted using the PureLink RNA Mini kit (ThermoFisher Scientific, USA) with digestion by DNAse (ThermoFisher Scientific, USA) into the columns.

### RNA sequencing

RNA integrity was performed using Bioanalyzer and libraries were prepared using a QuantSeq 3’ mRNA-Seq Library Prep Kit-FWD (Lexogen, Vienna, Austria) with 1μg RNA. The library concentration was measured by Qubit Fluorometer and Qubit dsDNA HS Assay Kit (Applied Biosystems), and the size distribution was determined using an Agilent D1000 ScreenTape System (Agilent Technologies). Sequencing was performed on the NextSeq 500 platform at the NGS facility core SELA. The sequencing data were aligned to the GRCh38 version of the human genome using STAR and the bamsort tool from biobambam2, for downstream processing of the BAM file, including merging, sorting, and marking of duplicates. We used featureCounts to count the number of reads that overlap each gene.

### Differential Expression Analysis

Raw RNA-seq counts were normalized using the median-of-ratios method implemented in DESeq2 ^15^. Technical replicates were collapsed into biological replicates using collapseReplicates and genes with fewer than 25 total counts across all samples were removed to eliminate low-expression noise. Normalized expression values were obtained using variance stabilizing transformation (VST) with blind = FALSE. PCA was performed on the transformed data to assess sample clustering.

The Likelihood Ratio Test (LRT) was employed to evaluate the effects of treatments and experimental factors. The full model included all experimental factors, while the reduced model included only the intercept. DEGs were identified using a false discovery rate (FDR) < 0.05 and an absolute log2 fold change > 0.58, unless otherwise mentioned. Exclusive DEGs for each treatment condition were determined by filtering genes uniquely responsive to a specific treatment while meeting these criteria. All steps were done in RStudio (v. 4.2.1). Graphs were generated using ggplot2. A total of 64 RNA-seq libraries were generated across six experimental conditions: control without light/photoactivation (n = 6), control with photoactivation (n = 6), rapamycin (n = 4), bafilomycin (n = 4), DMMB without photoactivation (n = 6), and DMMB with photoactivation (n = 6). All samples were included in downstream analyses, with no exclusions.

### Functional Enrichment Analysis

Significant DEGs were analyzed for enrichment in biological pathways and processes using clusterProfiler (GO terms) and KEGG pathway analysis. For clusterProfiler, pathways were considered significant if p-value < 0.01, with a minimum of two genes per pathway.

### Transcription Factor Prediction

To identify upstream transcriptional regulators, the CHEA3 method ^75^ was utilized to rank transcription factors related to DEGs according to the mean rank scoring system and log2FC change. Transcription factors exclusively associated with a single treatment were filtered and represented based on mean rank scoring and log2FC.

### Photosensitization protocol for Mitochondrial Respiration Analysis

Considering photoactivation response of the DMMB compound in cells, all experiments were carried out in four conditions: (*1*) cells without exposure to red light and in the absence of photosensitizer (*dark control*); (*2*) cells exposed only to red light, in the absence of photosensitizer (*Irradiated Control*); (3) cells incubated with photosensitizer and photoactivated with red light (*DMMB Irradiated*) and (4) cells incubated with photosensitizer in the absence of irradiation (*DMMB Dark*).

Twenty-four hours after sending, HaCaT cells were incubated with DMMB (10 nM) for one hour, then washed with PBS twice. Irradiation was carried out with specialized equipment (Ethik, Biolambda, Brazil) with a maximum emission wavelength at 630 nm. To provide 12 J cm−2, cells were irradiated for 9 min (23,5 mW). Immediately after the irradiation, 5 nM Bafilomycin-A1 (Bafilomycin) or 100 nM Rapamycin was added to the DMMB-irradiated groups in DMEM containing 1% FBS for twelve hours. Then the media was exchanged to DMEM containing 10% FBS. Cells were prepared for mitochondrial respiration 36h later. Then the media was exchanged to DMEM containing 10% FBS. Cells were prepared for mitochondrial respiration 36h later.

### Mitochondrial Respiration Analysis

The Seahorse XFe24 Analyzer (Agilent Technology, Santa Clara, CA, USA) was used for mitochondrial respiration analysis. The cells were washed three times with 500 μL DMEM/F-12 containing 1% P/S and 5 mM HEPES. The media did not contain bicarbonate or FBS. After incubation, plates were placed in the equipment and OCRs were measured under basal conditions, followed by different injections: 1) oligomycin, an ATP synthase inhibitor (complex V), and acting as OCR reducing agent; 2) carbonyl cyanide 3-chlorophenylhydrazone (CCCP), to induce maximum electron transport; 3) antimycin and rotenone (R/AA, final concentration, 1 μM each) complex III and complex I inhibitor, respectively.

Basal mitochondrial respiration was determined by monitoring OCR in the absence of any inhibitors and was calculated by reducing extracellular OCR through the inhibition of ATP synthase by oligomycin. The injection of oligomycin inhibits the complex V (F1–Fo ATP synthase), decreasing proton flux through complex V, and increasing the proton concentration, which reduces electron transport and oxygen consumption. The mitochondrial OCR remaining after oligomycin treatment is a measure of proton leak. The uncoupling agent, Carbonyl cyanide-4 (trifluoromethoxy) phenylhydrazone (FCCP), collapses the proton gradient and interferes with the mitochondrial membrane potential. As a consequence, electron flow through the electron transport chain is uninhibited, and it is possible to observe the maximal oxygen consumption by complex IV. The spare capacity was calculated by the difference between maximal and basal OCR. To conclude, all mitochondrial-associated respiration is eliminated by the addition of rotenone (a complex I inhibitor) and antimycin A (an inhibitor of cytochrome C reductase). A One-Way ANOVA followed by a Tukey post-test was used for each condition (Dark and irradiated). A p-value < 0.05 was considered significant (GraphPad Prism, v. 10). Each experimental group consisted of 4 to 5 independent biological replicates. The exception was the photoactivated DMMB group, which included only two replicates due to a technical error that prevented proper data acquisition from the remaining wells.

## DATA AVAILABILITY

Data is being deposited in the Gene Expression Omnibus (GEO) database. The processed data derived from the RNA sequencing are presented in Table S1. Additional data that support the findings in this manuscript can be obtained from the corresponding author upon request.

## ARTIFICIAL INTELLIGENCE STATEMENT

The authors used ChatGPT and Grammarly to improve readability and language, then reviewed and edited the content, taking full responsibility for the publication content.

## ACKNOWLEDGMENTS AND FUNDING

Baptista, M received grants from the São Paulo Research Foundation (2022/13066-9, 2021/08521-6, and 2013/07937-8). de Assis is supported by the Knut and Alice Wallenberg Foundation as a Wallenberg Molecular Medicine Fellow.

## AUTHOR’S CONTRIBUTION

Márcia Silvana Freire Franco: conceptualization, methodology, data curation, formal analysis, investigation, manuscript writing, and review. Felipe Gustavo Ravagnani: conceptualization, investigation, manuscript review. Suely Kazue Nagahashi Marie: methodology, data curation, formal analysis, manuscript review. Sueli Mieko Oba-Shinjo: methodology, data curation, manuscript review. Leonardo Vinicius Monteiro de Assis: conceptualization, methodology, data curation, formal analysis, supervision, manuscript writing and review. Maurício S. Baptista: conceptualization, methodology, funding, supervision, manuscript writing and review.

---

**Figure S1:**
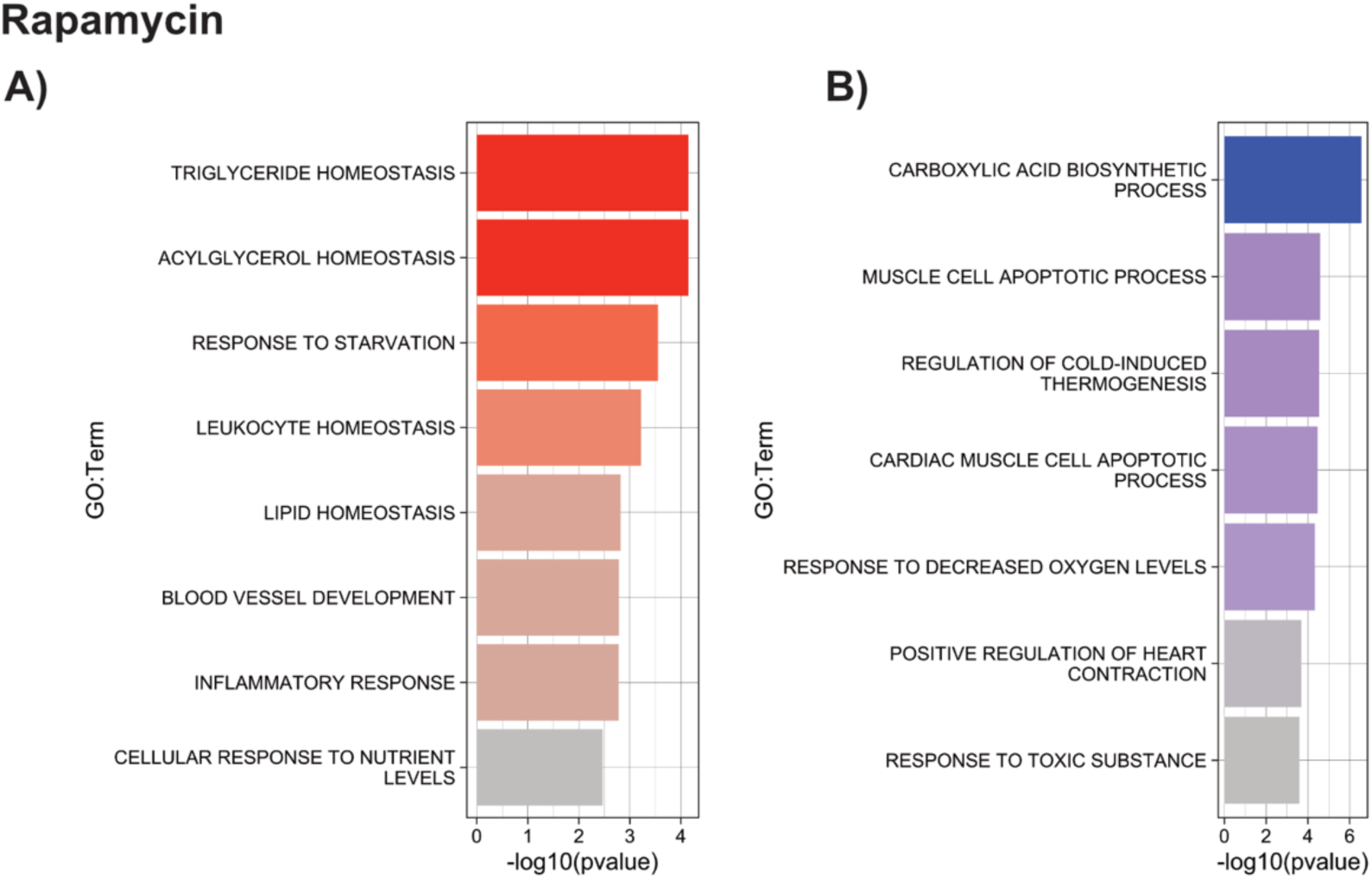
Enriched pathways for Rapamycin. A – B) Enrichment analyses conducted for upregulated (red) and downregulated genes (in blue) for Rapamycin-treated cells. Additional processes are shown in Table S1.

**Figure S2:**
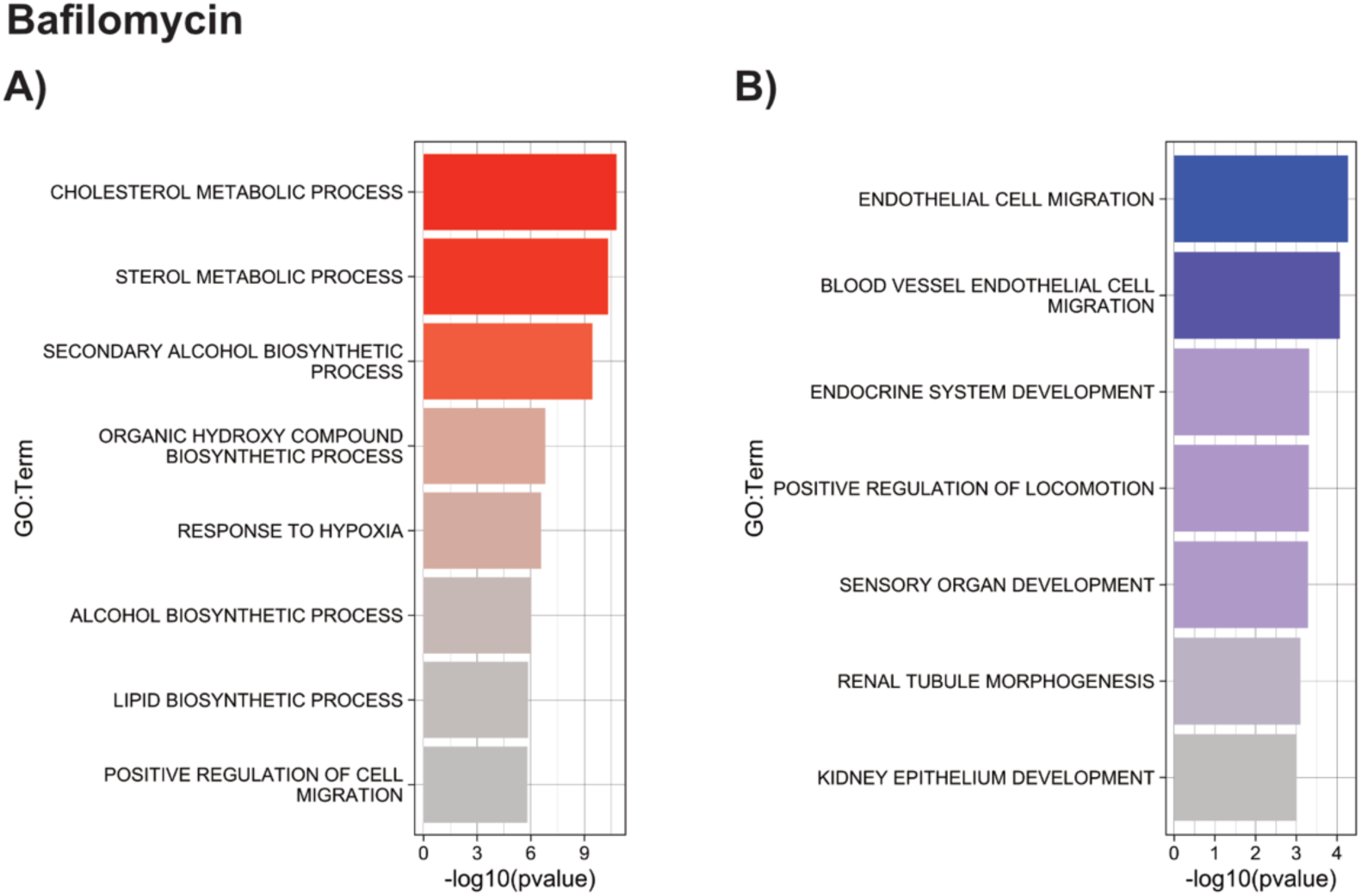
Enriched pathways for Bafilomycin. A – B) Enrichment analyses conducted for upregulated (red) and downregulated genes (in blue) for Bafilomycin-treated cells. Additional processes are shown in Table S1.

**Figure S3:**
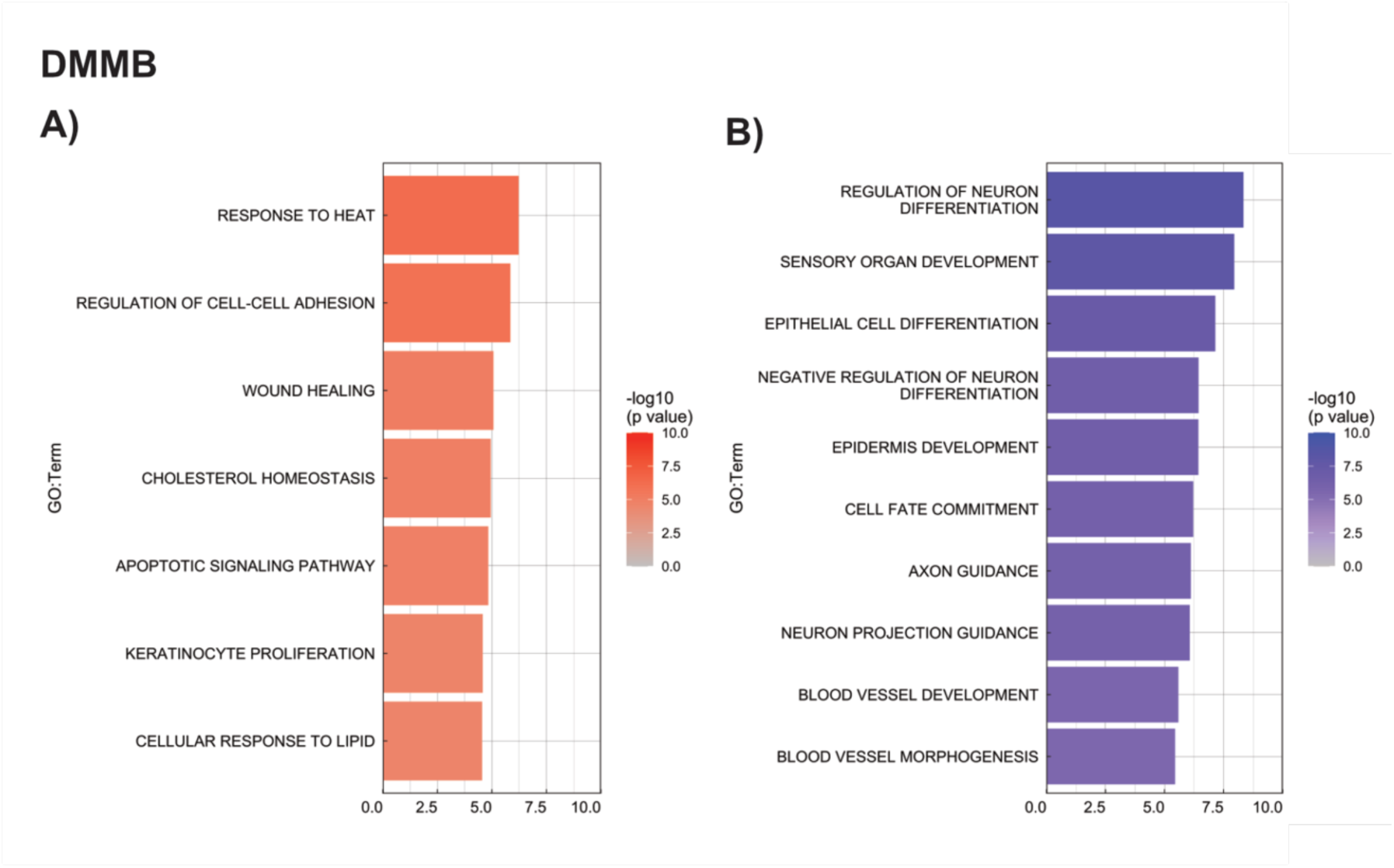
Enriched pathways for photoactivated DMMB. A – B) Enrichment analyses conducted for upregulated (red) and downregulated genes (in blue) for DMMB-treated cells. Additional processes are shown in Table S1.

## Notes

### Competing Interest Statement

The authors have declared no competing interest.

## REFERENCES

1. Feng Y, He D, Yao Z, Klionsky DJ. The machinery of macroautophagy. Cell Res 2014; 24:24–41.

2. Kroemer G, Levine B. Autophagic cell death: the story of a misnomer. Nat Rev Mol Cell Biol 2008; 9:1004–10.

3. Wang J, Zhu X, Zhang J, Wang H, Liu G, Bu Y, Yu J, Tian Y, Zhou H. AIE-Based Theranostic Agent: In Situ Tracking Mitophagy Prior to Late Apoptosis To Guide the Photodynamic Therapy. ACS Appl Mater Interfaces 2020; 12:1988–96.

4. Seiler A, Schneider M, Förster H, Roth S, Wirth EK, Culmsee C, Plesnila N, Kremmer E, Rådmark O, Wurst W, et al. Glutathione peroxidase 4 senses and translates oxidative stress into 12/15-lipoxygenase dependent– and AIF-mediated cell death. Cell Metab 2008; 8:237–48.

5. Shackley DC, Haylett A, Whitehurst C, Betts CD, O’Flynn K, Clarke NW, Moore JV. Comparison of the cellular molecular stress responses after treatments used in bladder cancer. BJU Int 2002; 90:924–32.

6. Duan X, Chen B, Cui Y, Zhou L, Wu C, Yang Z, Wen Y, Miao X, Li Q, Xiong L, et al. Ready player one? Autophagy shapes resistance to photodynamic therapy in cancers. Apoptosis Int J Program Cell Death 2018; 23:587–606.

7. Roberts DJ, Tan-Sah VP, Ding EY, Smith JM, Miyamoto S. Hexokinase-II positively regulates glucose starvation-induced autophagy through TORC1 inhibition. Mol Cell 2014; 53:521–33.

8. Wei X, Manandhar L, Kim H, Chhetri A, Hwang J, Jang G, Park C, Park R. Pexophagy and Oxidative Stress: Focus on Peroxisomal Proteins and Reactive Oxygen Species (ROS) Signaling Pathways. Antioxid Basel Switz 2025; 14:126.

9. Martins WK, Santos NF, Rocha C de S, Bacellar IOL, Tsubone TM, Viotto AC, Matsukuma AY, Abrantes AB de P, Siani P, Dias LG, et al. Parallel damage in mitochondria and lysosomes is an efficient way to photoinduce cell death. Autophagy 2019; 15:259–79.

10. Min DH, Kim D, Hong ST, Kim J, Kim MJ, Kwon S-H, Kim A, Lee J-Y. Bafilomycin A1 induces colon cancer cell death through impairment of the endolysosome system dependent on iron. Sci Rep 2025; 15:5148.

11. Zhdanov AV, Dmitriev RI, Papkovsky DB. Bafilomycin A1 activates respiration of neuronal cells via uncoupling associated with flickering depolarization of mitochondria. Cell Mol Life Sci CMLS 2011; 68:903–17.

12. Kennedy BK, Lamming DW. The Mechanistic Target of Rapamycin: The Grand ConducTOR of Metabolism and Aging. Cell Metab 2016; 23:990–1003.

13. He L, Zhang J, Zhao J, Ma N, Kim SW, Qiao S, Ma X. Autophagy: The Last Defense against Cellular Nutritional Stress. Adv Nutr Bethesda Md 2018; 9:493–504.

14. Mauvezin C, Neufeld TP. Bafilomycin A1 disrupts autophagic flux by inhibiting both V-ATPase-dependent acidification and Ca-P60A/SERCA-dependent autophagosome-lysosome fusion. Autophagy 2015; 11:1437–8.

15. Love MI, Huber W, Anders S. Moderated estimation of fold change and dispersion for RNA-seq data with DESeq2. Genome Biol 2014; 15:550.

16. Meijer AJ, Lorin S, Blommaart EF, Codogno P. Regulation of autophagy by amino acids and MTOR-dependent signal transduction. Amino Acids 2015; 47:2037–63.

17. Panwar V, Singh A, Bhatt M, Tonk RK, Azizov S, Raza AS, Sengupta S, Kumar D, Garg M. Multifaceted role of mTOR (mammalian target of rapamycin) signaling pathway in human health and disease. Signal Transduct Target Ther 2023; 8:375.

18. Sarkar S. Regulation of autophagy by mTOR-dependent and mTOR-independent pathways: autophagy dysfunction in neurodegenerative diseases and therapeutic application of autophagy enhancers. Biochem Soc Trans 2013; 41:1103–30.

19. Redmann M, Benavides GA, Berryhill TF, Wani WY, Ouyang X, Johnson MS, Ravi S, Barnes S, Darley-Usmar VM, Zhang J. Inhibition of autophagy with bafilomycin and chloroquine decreases mitochondrial quality and bioenergetic function in primary neurons. Redox Biol 2016; 11:73–81.

20. Garza-Lombó C, Schroder A, Reyes-Reyes EM, Franco R. mTOR/AMPK signaling in the brain: Cell metabolism, proteostasis and survival. Curr Opin Toxicol 2018; 8:102–10.

21. Birnbaum MJ. Activating AMP-activated protein kinase without AMP. Mol Cell 2005; 19:289–90.

22. Meley D, Bauvy C, Houben-Weerts JHPM, Dubbelhuis PF, Helmond MTJ, Codogno P, Meijer AJ. AMP-activated protein kinase and the regulation of autophagic proteolysis. J Biol Chem 2006; 281:34870–9.

23. Sabatini DM. mTOR and cancer: insights into a complex relationship. Nat Rev Cancer 2006; 6:729– 34.

24. Hardie DG. The AMP-activated protein kinase pathway--new players upstream and downstream. J Cell Sci 2004; 117:5479–87.

25. Liu J, Zhang H. Zinc Finger and BTB Domain-Containing 20: A Newly Emerging Player in Pathogenesis and Development of Human Cancers. Biomolecules 2024; 14:192.

26. Chakrabarti P, English T, Shi J, Smas CM, Kandror KV. Mammalian target of rapamycin complex 1 suppresses lipolysis, stimulates lipogenesis, and promotes fat storage. Diabetes 2010; 59:775–81.

27. Zheng Y, Neculai D, Fairn GD. S-acylation of PNPLA2/ATGL: a necessity for triacylglycerol lipolysis and lipophagy in hepatocytes. Autophagy 2025; 21:494–6.

28. Kryszczuk M, Kowalczuk O. Significance of NRF2 in physiological and pathological conditions an comprehensive review. Arch Biochem Biophys 2022; 730:109417.

29. Gong SS, Guerrini L, Basilico C. Regulation of asparagine synthetase gene expression by amino acid starvation. Mol Cell Biol 1991; 11:6059–66.

30. Krall AS, Mullen PJ, Surjono F, Momcilovic M, Schmid EW, Halbrook CJ, Thambundit A, Mittelman SD, Lyssiotis CA, Shackelford DB, et al. Asparagine couples mitochondrial respiration to ATF4 activity and tumor growth. Cell Metab 2021; 33:1013–1026.e6.

31. Pavlova NN, Hui S, Ghergurovich JM, Fan J, Intlekofer AM, White RM, Rabinowitz JD, Thompson CB, Zhang J. As Extracellular Glutamine Levels Decline, Asparagine Becomes an Essential Amino Acid. Cell Metab 2018; 27:428–438.e5.

32. Chiodi I, Perini C, Berardi D, Mondello C. Asparagine sustains cellular proliferation and c-Myc expression in glutamine-starved cancer cells. Oncol Rep 2021; 45:96.

33. Wang R, Yu Z, Sunchu B, Shoaf J, Dang I, Zhao S, Caples K, Bradley L, Beaver LM, Ho E, et al. Rapamycin inhibits the secretory phenotype of senescent cells by a Nrf2-independent mechanism. Aging Cell 2017; 16:564–74.

34. Xu Y, Tai W, Qu X, Wu W, Li Z, Deng S, Vongphouttha C, Dong Z. Rapamycin protects against paraquat-induced pulmonary fibrosis: Activation of Nrf2 signaling pathway. Biochem Biophys Res Commun 2017; 490:535–40.

35. Liao M, Yao D, Wu L, Luo C, Wang Z, Zhang J, Liu B. Targeting the Warburg effect: A revisited perspective from molecular mechanisms to traditional and innovative therapeutic strategies in cancer. Acta Pharm Sin B 2024; 14:953–1008.

36. Deng X, Wang Q, Cheng M, Chen Y, Yan X, Guo R, Sun L, Li Y, Liu Y. Pyruvate dehydrogenase kinase 1 interferes with glucose metabolism reprogramming and mitochondrial quality control to aggravate stress damage in cancer. J Cancer 2020; 11:962–73.

37. Cao W, Zeng Z, Pan R, Wu H, Zhang X, Chen H, Nie Y, Yu Z, Lei S. Hypoxia-Related Gene FUT11 Promotes Pancreatic Cancer Progression by Maintaining the Stability of PDK1. Front Oncol 2021; 11:675991.

38. Kim J, Tchernyshyov I, Semenza GL, Dang CV. HIF-1-mediated expression of pyruvate dehydrogenase kinase: a metabolic switch required for cellular adaptation to hypoxia. Cell Metab 2006; 3:177–85.

39. Naquet P, Kerr EW, Vickers SD, Leonardi R. Regulation of coenzyme A levels by degradation: the “Ins and Outs.” Prog Lipid Res 2020; 78:101028.

40. Feng X, Zhang L, Xu S, Shen A-Z. ATP-citrate lyase (ACLY) in lipid metabolism and atherosclerosis: An updated review. Prog Lipid Res 2020; 77:101006.

41. Gu J, Zhu N, Li H-F, Zhao T-J, Zhang C-J, Liao D-F, Qin L. Cholesterol homeostasis and cancer: a new perspective on the low-density lipoprotein receptor. Cell Oncol Dordr Neth 2022; 45:709–28.

42. Ha NT, Lee CH. Roles of Farnesyl-Diphosphate Farnesyltransferase 1 in Tumour and Tumour Microenvironments. Cells 2020; 9:2352.

43. Hu L-D, Wang J, Chen X-J, Yan Y-B. Lanosterol modulates proteostasis via dissolving cytosolic sequestosomes/aggresome-like induced structures. Biochim Biophys Acta BBA – Mol Cell Res 2020; 1867:118617.

44. Miyazaki S, Shimizu N, Miyahara H, Teranishi H, Umeda R, Yano S, Shimada T, Shiraishi H, Komiya K, Katoh A, et al. DHCR7 links cholesterol synthesis with neuronal development and axonal integrity. Biochem Biophys Res Commun 2024; 712–713:149932.

45. Porter FD, Herman GE. Malformation syndromes caused by disorders of cholesterol synthesis. J Lipid Res 2011; 52:6–34.

46. Mao Z, Feng M, Li Z, Zhou M, Xu L, Pan K, Wang S, Su W, Zhang W. ETV5 Regulates Hepatic Fatty Acid Metabolism Through PPAR Signaling Pathway. Diabetes 2021; 70:214–26.

47. Puli OR, Danysh BP, McBeath E, Sinha DK, Hoang NM, Powell RT, Danysh HE, Cabanillas ME, Cote GJ, Hofmann M-C. The Transcription Factor ETV5 Mediates BRAFV600E-Induced Proliferation and TWIST1 Expression in Papillary Thyroid Cancer Cells. Neoplasia N Y N 2018; 20:1121–34.

48. Weng L, Cheng Z, Qiu Z, Shi J, Chen L, He C, Wang L, Jin F. Integration of bioinformatics analysis reveals ZNF248 as a potential prognostic and immunotherapeutic biomarker for LIHC: machine learning and experimental evidence. Am J Cancer Res 2024; 14:5230–50.

49. Liu Z, Kruhlak MJ, Thiele CJ. Zinc finger transcription factor CASZ1b is involved in the DNA damage response in live cells. Biochem Biophys Res Commun 2023; 663:171–8.

50. Droll SH, Zhang BJ, Levine MC, Xue C, Ho PJ, Bao X. CASZ1 Is Essential for Skin Epidermal Terminal Differentiation. J Invest Dermatol 2024; 144:2029–38.

51. Webster CP, Smith EF, Bauer CS, Moller A, Hautbergue GM, Ferraiuolo L, Myszczynska MA, Higginbottom A, Walsh MJ, Whitworth AJ, et al. The C9orf72 protein interacts with Rab1a and the ULK1 complex to regulate initiation of autophagy. EMBO J 2016; 35:1656–76.

52. Pietrement C, Sun-Wada G-H, Silva ND, McKee M, Marshansky V, Brown D, Futai M, Breton S. Distinct expression patterns of different subunit isoforms of the V-ATPase in the rat epididymis. Biol Reprod 2006; 74:185–94.

53. Reid E, Connell J, Edwards TL, Duley S, Brown SE, Sanderson CM. The hereditary spastic paraplegia protein spastin interacts with the ESCRT-III complex-associated endosomal protein CHMP1B. Hum Mol Genet 2005; 14:19–38.

54. Takahashi Y, He H, Tang Z, Hattori T, Liu Y, Young MM, Serfass JM, Chen L, Gebru M, Chen C, et al. An autophagy assay reveals the ESCRT-III component CHMP2A as a regulator of phagophore closure. Nat Commun 2018; 9:2855.

55. Xia Q, Liu X, Zhong L, Qu J, Dong L. SMURF1 mediates damaged lysosomal homeostasis by ubiquitinating PPP3CB to promote the activation of TFEB. Autophagy 2025; 21:530–47.

56. Papadopoulos C, Kirchner P, Bug M, Grum D, Koerver L, Schulze N, Poehler R, Dressler A, Fengler S, Arhzaouy K, et al. VCP/p97 cooperates with YOD1, UBXD1 and PLAA to drive clearance of ruptured lysosomes by autophagy. EMBO J 2017; 36:135–50.

57. Jiang B, Zhao Y, Shi M, Song L, Wang Q, Qin Q, Song X, Wu S, Fang Z, Liu X. DNAJB6 Promotes Ferroptosis in Esophageal Squamous Cell Carcinoma. Dig Dis Sci 2020; 65:1999–2008.

58. Wang Z, Wu S, Zhu C, Shen J. The role of ferroptosis in esophageal cancer. Cancer Cell Int 2022; 22:266.

59. Zou Y, Xiong J, Ma K, Wang A-Z, Qian K-J. Rac2 deficiency attenuates CCl4-induced liver injury through suppressing inflammation and oxidative stress. Biomed Pharmacother 2017; 94:140–9.

60. Krejsa CM, Franklin CC, White CC, Ledbetter JA, Schieven GL, Kavanagh TJ. Rapid activation of glutamate cysteine ligase following oxidative stress. J Biol Chem 2010; 285:16116–24.

61. Pickrell AM, Youle RJ. The Roles of PINK1, Parkin, and Mitochondrial Fidelity in Parkinson’s Disease. Neuron 2015; 85:257–73.

62. Soto-Avellaneda A, Morrison BE. Signaling and other functions of lipids in autophagy: a review. Lipids Health Dis 2020; 19:214.

63. Xie Y, Li J, Kang R, Tang D. Interplay Between Lipid Metabolism and Autophagy. Front Cell Dev Biol 2020; 8:431.

64. Ravnskjaer K, Frigerio F, Boergesen M, Nielsen T, Maechler P, Mandrup S. PPARdelta is a fatty acid sensor that enhances mitochondrial oxidation in insulin-secreting cells and protects against fatty acid-induced dysfunction. J Lipid Res 2010; 51:1370–9.

65. de Assis LVM, Oster H. The circadian clock and metabolic homeostasis: entangled networks. Cell Mol Life Sci 2021; 78:4563–87.

66. Sato T, Greco CM. Expanding the link between circadian rhythms and redox metabolism of epigenetic control. Free Radic Biol Med 2021; 170:50–8.

67. Wilking M, Ndiaye M, Mukhtar H, Ahmad N. Circadian rhythm connections to oxidative stress: implications for human health. Antioxid Redox Signal 2013; 19:192–208.

68. Pickles S, Vigié P, Youle RJ. The art of mitochondrial maintenance. Curr Biol CB 2018; 28:R170– 85.

69. Whitworth AJ, Pallanck LJ. The PINK1/Parkin pathway: a mitochondrial quality control system? J Bioenerg Biomembr 2009; 41:499–503.

70. Chen Y, Lam TT, Yu X, Vasiliou V. 6 – Differential Redox Changes in Liver Mitochondrial Proteomes from Phenotypically Distinct Mouse Models of Glutathione Deficiency. Free Radic Biol Med 2017; 112:22.

71. Di Noia MA, Todisco S, Cirigliano A, Rinaldi T, Agrimi G, Iacobazzi V, Palmieri F. The human SLC25A33 and SLC25A36 genes of solute carrier family 25 encode two mitochondrial pyrimidine nucleotide transporters. J Biol Chem 2014; 289:33137–48.

72. von Bohlen Und Halbach O. Controlling glutathione entry into mitochondria: potential roles for SLC25A39 in health and (treatment of) disease. Signal Transduct Target Ther 2022; 7:75.

73. de Assis LVM, Tonolli PN, Moraes MN, Baptista MS, de Lauro Castrucci AM. How does the skin sense sun light? An integrative view of light sensing molecules. J Photochem Photobiol C Photochem Rev 2021; 47:100403.

74. Tonolli PN, Marie SKN, Oba-Shinjo SM, de Assis LVM, Baptista MS. Stage-specific phenotypic and transcriptional alterations in HaCaT keratinocytes exposed to acute and chronic blue light. Photochem Photobiol 2025;

75. Keenan AB, Torre D, Lachmann A, Leong AK, Wojciechowicz ML, Utti V, Jagodnik KM, Kropiwnicki E, Wang Z, Ma’ayan A. ChEA3: transcription factor enrichment analysis by orthogonal omics integration. Nucleic Acids Res 2019; 47:W212–24.

